# Organized spatial patterns of activated β_2_ integrins in arresting neutrophils

**DOI:** 10.1101/323279

**Authors:** Zhichao Fan, William Bill Kiosses, Dirk M. Zajonc, M. Amin Arnaout, Edgar Gutierrez, Alex Groisman, Klaus Ley

## Abstract

The transition from leukocyte rolling to firm adhesion is called arrest. β_2_ integrins are required for neutrophil arrest^1^. Chemokines can trigger neutrophil arrest in vivo^2^ and in vitro^3^. Resting integrins^4^ exist in a “bent-closed” conformation, i.e., not extended (E^−^) and not high affinity (H^−^), unable to bind ligand. Electron microscopic images of isolated β_2_ integrins in “open” and “closed” conformations^5^ inspired the switchblade model of integrin activation from E^−^H^−^ to E^+^H^−^ to E^+^H^+6^^7^. Recently^8^, we discovered an alternative pathway of integrin activation from E^−^H^−^ to E^−^H^+^ to E^+^H^+^. Spatial patterning of activated integrins is thought to be required for effective arrest, but so far only diffraction-limited localization maps of activated integrins exist^8^. Here, we combine superresolution microscopy with molecular modeling to identify the molecular patterns of H^+^E^−^, H^−^E^+^, and H^+^E^+^ activated integrins on primary human neutrophils. At the time of neutrophil arrest, E^+^H^+^ integrins form oriented (non-random) nanoclusters that contain a total of 4,625±369 E^+^H^+^ β_2_ integrin molecules.

---

The switchblade model posits that agonist-triggered cell activation (“inside-out”) first triggers extension (E^−^→E^+^) of both the α and the β extracellular domains of integrins, which can then assume the high-affinity conformation (H^−^→H^+^) of the ligand binding site located in the I (also known as A) domain of the α subunit. Thus, the switchblade model posits a sequence of E^−^H^−^→E^+^H^−^→E^+^H^+^. In a recent study using quantitative dynamic footprinting (qDF) microscopy, a combination of a microfluidic flow chamber with variable angle total internal reflection fluorescence (TIRF) microscopy^9^, we discovered the E^−^H^−^→E^−^H^+^→E^+^H^+^ pathway and showed that activated β_2_ integrins are clustered on the neutrophil surface even prior to chemokine activation, i.e., during rolling^8^. We used two monoclonal antibodies (mAbs) that report the β_2_ integrin conformation: mAb KIM127^10^ recognizes an epitope in the knee of β_2_ that is not accessible when the knee is bent and thus reports extension (E^+^)^11^ and mAb24^12^ recognizes an epitope in the β_2_ I-like domain that appears when the α I/A domain assumes the high affinity (H^+^) conformation^13^. Neither antibody interferes with ligand binding, and KIM127 and mAb24 do not cross-block each other^8^. Rolling on P-selectin induced clusters of KIM127^+^ E^+^H^−^ β_2_ integrins. Exposure to the chemokine interleukin 8 (IL-8) induced mAb24 binding. Unexpectedly, we found activated E^−^H^+^ (high-affinity bent) clusters that were stabilized by binding to the β_2_ integrin ligands ICAM-1 and ICAM-3 in cis. By this cis binding, E-H+ integrins exert a strong anti-adhesion effect on both human and mouse neutrophils. Because qDF is diffraction-limited, it (1) remained unclear whether apparent E^+^H^+^ clusters identified by qDF contain only E^+^H^+^ integrins or a mixture of E^+^H^−^ and E^−^H^+^ integrins and (2) whether ordered patterns exist in activated integrin arrangement on the cell surface. Patterned integrin arrays have been suggested to be important for integrin function^14^, but have never been observed in cells.

Recently, super-resolution microscopy techniques have been developed^15,16^ and applied to integrin biology^17^. Springer et al. studied migrating, not rolling and arresting cells, using a cell line (Jurkat), not primary cells. There was no flow, no estimate of the force per integrin and no mapping of different integrin conformations over the footprint of the cell. Although this study was informative in supporting integrin extension by direct superresolution measurements, it did not address patterns of integrins on the cell surface.

To study β_2_ integrin activation in primary human neutrophils during arrest with super-resolution microscopy, we developed a strategy to fix the cells exactly at the time of their IL-8-triggered arrest (Extended Data Fig. 1a-c). Comparing qDF-based images of the footprint of arrested neutrophils with TIRF images of the same cells after fixation shows that there is no significant change in the morphology of the cell footprint after fixation (Extended Data Fig. 1d and e). Conventional TIRF microscopy of the neutrophil footprints at the time of arrest showed maps (Fig. 1a) of activated integrins labeled with KIM127 Fab fragments (indicating E^+^) and mAb24 Fab fragments (indicating H^+^), similar to what we reported previously using qDF live cell imaging^8^. Processing the TIRF images using a smart segmentation algorithm^8^ suggests large clusters of E^−^H^+^ (cyan), E^+^H^−^ (magenta) and E^+^H^+^ (white) integrins (Fig. 1b). The similarity of the integrin clusters obtained by TIRF and qDF^18^ suggests that (1) fixation did not introduce major changes in the distribution and expression of activated β_2_ integrins and (2) Fab fragments were similar to intact antibodies in their ability to label activated β_2_ integrins. Next, we applied stochastic optical reconstruction microscopy (STORM)^15^ to achieve higher resolution maps of activated β_2_ integrins in the footprint of arrested neutrophils (Fig. 1c). Comparing Fig. 1c and 1a shows that, as expected, a much higher resolution was indeed achieved by STORM.

**Fig. 1.**
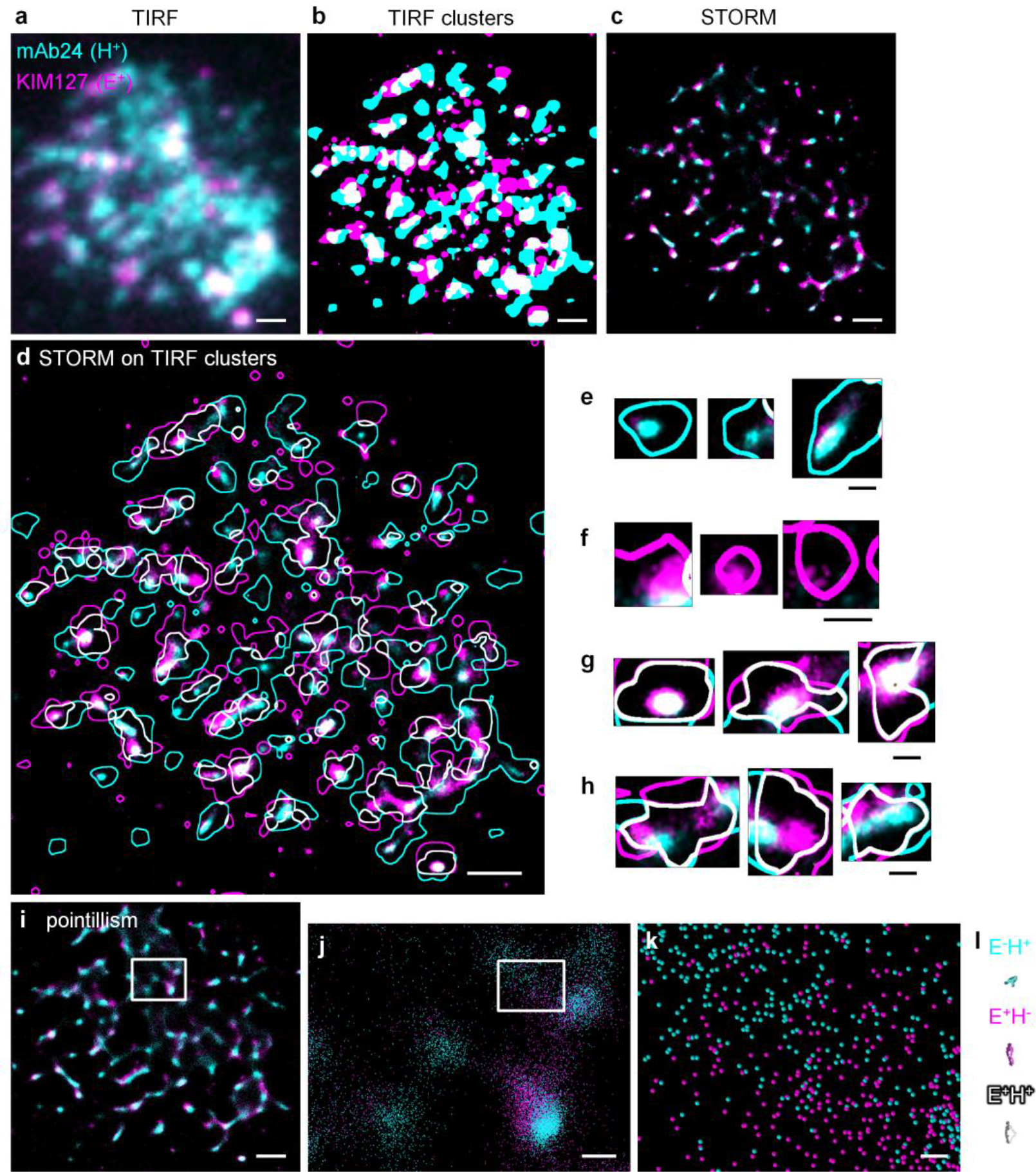
Super-resolution STORM imaging of β_2_ integrin activation on arrested human neutrophils. **a** Primary human neutrophils rolling on P-selectin and ICAM-1 at a wall shear stress of 6 dyn/cm^2^ were exposed to IL-8 and Fab fragments of the β_2_ integrin extension reporter KIM127 (E^+^, magenta) and the high-affinity reporter mAb24 (H^+^, cyan), immediately fixed and imaged by TIRF (entire TIRF footprint, raw image shown). **b** Binary image of **a** using smart segmentation as in^8^ **c** STORM buffer was introduced and blinking events recorded for 10,000 frames per channel over 10 minutes, corrected for stochastic motion and drift as in supplementary Methods to obtain a raw super-resolution STORM image of the footprint of arrested human neutrophils. **d**-**h** STORM image overlaid with the outlines of binary TIRF clusters (from **b**). Zoomed-in examples of E^−^H^+^ (cyan, **e**) and E^+^H^−^ TIRF clusters (magenta, **f**). Some white E^+^H^+^ TIRF clusters showed true colocalization of KIM127 and mAb24 in STORM (**g**, white). **h** shows examples of clusters that appeared colocalized in TIRF but were truly composed of E^−^H^+^ and E^+^H^−^ areas as revealed by STORM. **i**-**k** Pointillism map of locations of H^+^ and E^+^ generated from STORM imaging (details in Supplemental Methods), zoomed-in images in **j** (box in **i**) and **k** (box in **j**). **l** Structures of three active conformations of β_2_ integrins (E^−^H^+^ cyan, E^+^H^−^ magenta, E^+^H^+^ gray) at the same scale of **k**. Scale bars are 1μm for **a**-**d** and **i**, 200nm for **e**-**h** and **j**, and 30nm for **k**.

From STORM data, pointillism maps (Fig. 1i-k) can be constructed^19^, where each point represents the most likely location of the peak of a 2-dimensional Gaussian distribution of photons emanating from one fluorochrome called “blinks”. The location of the most likely position of the peaks of the Gaussians for KIM127 and mAb24 approach the molecular scale of β_2_ integrins (Fig. 1l). The resolution of our STORM setup (localization precision, σ) was around 15 nm (Extended Data Fig. 2a). Although this is much better than the ~250 nm resolution of the qDF or TIRF images, it is still not sufficient to resolve individual integrin molecules (size ~20×10×5 nm) with confidence.

The previous qDF-based work had suggested three types of clusters of integrins, enriched in E^−^H^+^, E^+^H^−^ or E^+^H^+^. We reasoned that E^−^H^+^ clusters mainly contained E^−^H^+^ integrins and E^+^H^−^ clusters mainly contained E^+^H^−^ integrins (Extended Data Fig. 3). This was indeed confirmed by STORM microscopy (Fig. 1d-f). We next hypothesized that E^+^H^+^ clusters identified by qDF microscopy might be of two types, either containing mainly E^+^H^+^ β_2_ integrins or a mixture of E^+^H^−^ and E^−^H^+^ (and perhaps E^+^H^+^) integrins (Extended Data Fig. 3). However, qDF had insufficient resolution to determine the true molecular composition of clusters. Analyzing the content of E^+^H^+^ TIRF clusters (Fig. 1d, g-h) revealed that indeed both types of clusters existed: some were dominated by E^+^H^+^ β_2_ integrins (Fig. 1g), and others contained a mixture of E^+^H^−^ and E^−^H^+^ integrins (Fig. 1h).

To push the resolution beyond the limits of the current STORM algorithm, we took advantage of the knowledge of the integrin shapes, thus allowing us to combine molecular modeling with STORM microscopy in a procedure we call Super-STORM. First, we filtered the data for areas in which integrins could physically bind ligands in trans. Since the total length of an extended β_2_ integrin bound to ICAM-1 is ~50 nm^8,17^, and microvilli on the surface of rolling neutrophils are up to ~200 nm high^20^, only integrins on the “tops” of microvilli can reach (and bind) ICAM-1, and other integrins on the “sides” of the microvilli or in the “valleys” between the microvilli are too far away from the adhesive substrate to contribute to binding^8^, even though such integrin molecules may be in the E^+^H^+^ conformation (Extended Data Fig. 4a). Since this study is focused on neutrophil arrest, and since only integrin molecules anchored in regions of the plasma membrane that were less than 50 nm away from the coverslip, we focused on these integrin molecules. We took advantage of the fact that the intensity of the TIRF signal emanating from the homogeneous membrane label (CD16, a glycosylphosphatidylinositol-anchored, GPI-anchored protein) mono-exponentially decays with the distance from the substrate^8,18,21,22^. From the membrane TIRF signal, we constructed a three-dimensional (3D) “hills and valleys” plot and gated it for a distance of 50 nm from the coverslip or less (Fig. 2a, Extended Data Fig. 4). Applying this 50 nm “height” gate to the entire footprint of a neutrophil revealed that only ~11% of the neutrophil footprint was within 50 nm of the coverslip (Fig 2b-d). Visual comparison of Fig 2c and d illustrates that indeed the vast majority of activated β_2_ integrins of all three conformations, E^+^H^−^, E^−^H^+^ and E^+^H^+^, are outside the reach of ICAM-1 in cis and thus irrelevant to arrest. Quantification shows that ~25% H^+^ and ~23% E^+^ β_2_ integrin molecules (Fig. 2e and f) are within reach of the ICAM-1 substrate.

**Fig. 2.**
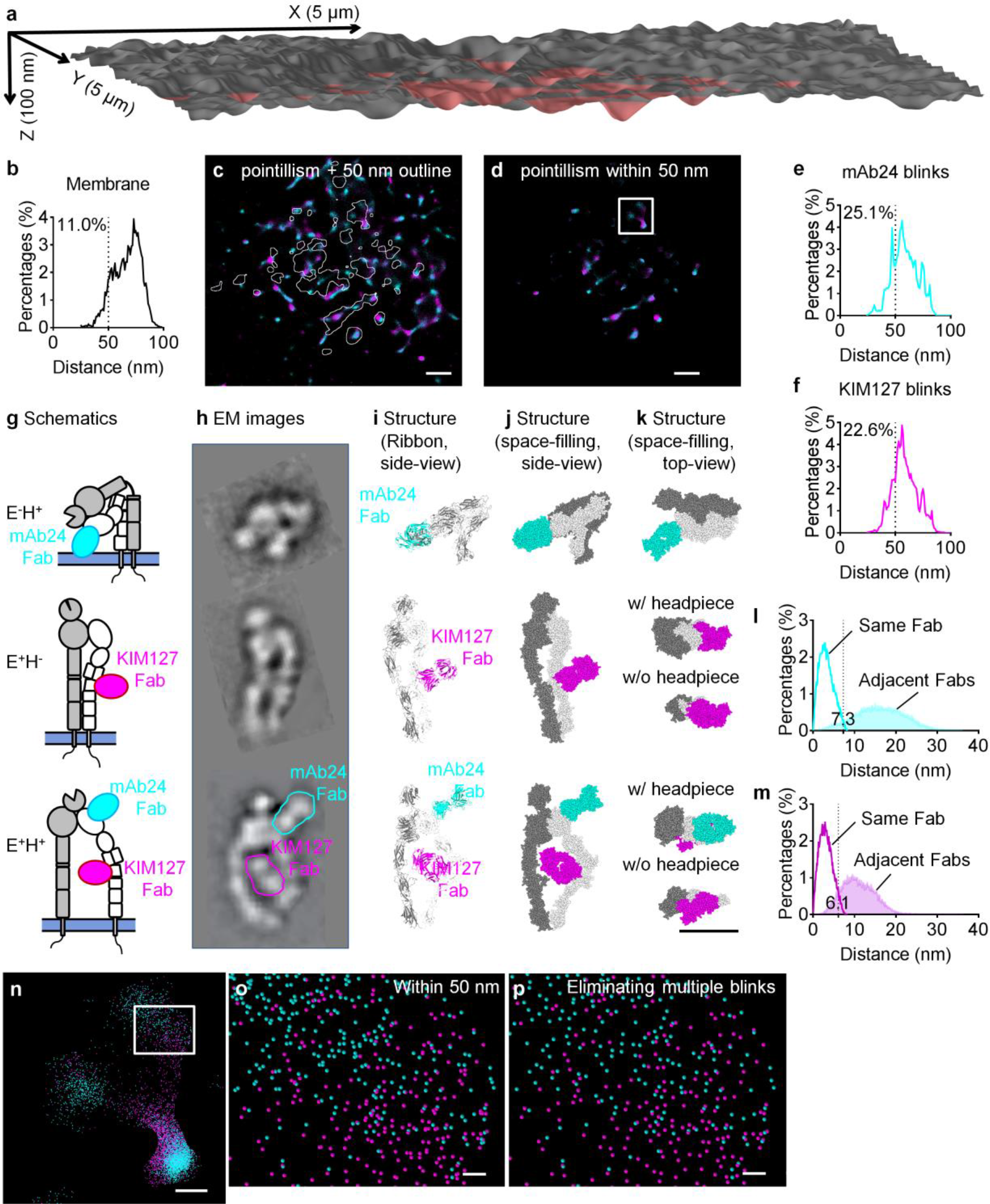
Molecular modeling for improved imaging processing (Super-STORM). **a** 3D topography of a neutrophil footprint generated from the TIRF image of anti-CD16-Alexa Fluor 488 (AF488) as in^8,18,21^. CD16 is a homogeneously distributed GPI-anchored protein. Area within 50 nm to the substrate highlighted in red. X, Y, and Z scales indicated. **b** Distribution of distances between the neutrophil membrane and the coverslip substrate. **c** STORM pointillism map overlaid with white outlines showing the area within 50 nm of the substrate. **d** Pointillism map as in c), gated for areas within 50 nm of the substrate. **e**-**f** Distribution of distances between mAb24 blinks (**e**) and KIM127 blinks (**f**) within 50 nm of the substrate. **g** Side-view schematics of E^−^H^+^, E^+^H^−^ and E^+^H^+^ β_2_ integrins with KIM127 and mAb24 Fabs bound, **h** EM images, **i** ribbon, **j** space-filling. **k** space-filling structure, top view, see also Supplementary Movies 1-3. EM images were adapted from^24,25^. The Fab of CBR LFA-1/2 in the EM images of E^+^H^−^ and E^+^H^+^ and the ligand C3c in the EM image of E^+^H^+^ were removed with photoshop for clarity (Extended Data Fig. 3). E^−^H^+^ β_2_ integrin is from the published crystal structure (PDB: 4NEH)^27^; E^+^H^−^ is modeled by unfolding the headpiece of E^−^H^−^ (PDB: 3K6S)^26^ to extension. For the E^+^H^+^ modeling, the hybrid domain swing out is superposed with PDB: 3FCU^28^. α-chain in grey and β-chain in white. mAb24 and KIM127 Fab were docked to their binding sites^11,13^. **l**-**m** Distance distributions of localizations emanating from the same Fab (open) or from Fabs bound to adjacent integrin molecules (filled) of mAb24 (**l**) or KIM127 (**m**), based on simulations of 64,0 randomly oriented integrin molecules. **n**-**p** Zoomed-in pointillism maps of **d**. **n** (box in **d**) and **o** (box in **n**). **p** The pointillism map after removing multiple blinks from **o**, based on the cutoffs indicated in l and m. The fields-of-view in **o**-**p** are the same as in Fig. 1k. Scale bars are 1 μm for **c**-**d**, 10nm for **i**-**k**, 200nm for **n**, and 30nm for **o**-**p**.

Next, we concentrated our attention on the orientation of these activated β_2_ integrins within 50 nm of the coverslip. Although the exact angle between an extended integrin and the membrane plane is not known^17^, a conservative assumption is that β_2_ integrin anchored in the plasma membrane at a distance of 50 nm from the coverslip must “stand up”, i.e. be oriented vertically, normal to the plasma membrane to bind ICAM-1 in trans. Thus, the microscopic image of such integrins is equivalent to the “top” view rather than a “side” view. We took advantage of the known structure of Fab fragments^23^ and applied molecular modeling to the Fab fragments bound to β_2_ integrins (Fig. 2g-k and Supplementary movies 1-3). Fig. 2g shows a schematic overview of the three β_2_ integrin conformations in side view, E^−^H^+^ (top), E^+^H^−^ (middle), and E^+^H^+^ (bottom). These schematics are based on rotary shadowing electron microscopy (EM) images (Fig. 2h, Extended Data Fig. 5)^24,25^. For clarity, the position of the mAb24 Fab is outlined in cyan and that of the KIM127 Fab in magenta. Since crystal structures of bent β_2_ integrins (E^−^H^− 26^ and E^−^H^+ 27^) and the headpiece for E^+^H^+^ integrin (with swung-out hybrid domain)^28^ are available (PDB: 3K6S, 4NEH, and 3FCU), and binding sites of mAb24^13^ and KIM127^11^ are known, we were able to construct ribbon diagrams (Fig. 2i) and space-filling models (Fig. 2j) of E^−^H^+^, E^+^H^−^ and E^+^H^+^ integrins. Because the shape of the three integrin conformations and the location of the bound Fab fragments is known, and only the Fab fragments are labeled, their possible locations are constrained by the projection of the integrin molecules in the “top” view (Fig. 2k).

In STORM, each fluorochrome can blink more than once^29^, and each Fab fragment may contain more than one fluorochrome. To accurately count molecules, multiple blinks emanating from within the projection of each Fab were merged and considered to represent the same Fab fragment (Extended Data Fig. 2b). We modeled the distribution of distances of blinks emanating from the same Fab and compared it to the distribution of distances emanating from Fab fragments bound to adjacent β_2_ integrin molecules (Fig. 2l-m). The distance distribution of blinks from Fabs bound to adjacent integrin molecules was obtained by modeling space filling adjacent β_2_ integrin (1,000 random simulations each of 64 adjacent conditions, Extended Data Fig. 6a-b). Both for mAb24 (Fig. 2l) and KIM127 (Fig. 2m), there was very little overlap between the two distributions (Supplemental Table 1). Thus, we used the crossing point to distinguish between photons emanating from the same Fab versus from Fabs bound to adjacent integrin molecules: mAb24 signals within 7.3 nm from each other and KIM127 signals within 6.1 nm from each other were assigned to be derived from the same Fab. The reason the projected area of the KIM127 Fab is slightly smaller is that the top-view of the E^+^H^−^ is smaller than that of the E^−^H^+^ β_2_ integrin (Fig. 2k). Filtering all blinks in the pointillism map by the proximity criteria of 7.3 nm for mAb24 and 6.1 nm for KIM127 reduced the number of blinks within 50 nm to the coverslip from ~35k and ~17k to ~11k and ~10k, respectively, (compare Fig. 2o and p). Since each Fab binds exactly one epitope in β_2_, this method allows us to count the number of E^+^ (KIM127^+^) and H^+^ (mAb24^+^) integrin molecules. This number was determined for each of 40 microvilli within 50 nm from the coverslip and showed 256±573 E^+^ and 292±648 H^+^ integrin molecules per microvillus (Supplemental Table 2).

To test whether we had successfully eliminated multiple blinks (which would result in an overestimation of activated integrin molecules), we compared the number of E^+^ (KIM127^+^) and H^+^ (mAb24^+^) STORM-based dots with the number of E^+^ and H^+^ integrins obtained by a completely independent method of counting the activated integrin molecules in the neutrophil footprint (Extended Data Fig. 7). As reported previously^8^, the number of molecules in the footprint can be determined by flow cytometry, using calibration beads with known numbers of antibody binding sites, and expressing the footprint area as a percentage of total cell surface area. The number of mAb24 and KIM127 epitopes within 50 nm to the coverslip was estimated based on the surface maps^8^ and yielded an average of ~12k mAb24 and ~21k KIM127 epitopes within 50 nm per footprint, similar to ~12k mAb24 and ~10k KIM127 epitopes within 50 nm obtained by direct molecule counting from STORM and molecular modeling.

Neutrophil arrest is caused by E^+^H^+^ β_2_ integrins binding ICAM-1 in trans^8,30^. The E^+^H^+^ integrin molecules within “reach” of ICAM-1 (membrane within 50 nm from the coverslip) integrins have both KIM127 and mAb24 Fabs bound. If a KIM127 Fab and a mAb24 Fab is bound to the same integrin molecule, we call this a “pair” and consider that integrin to be in the E^+^H^+^ conformation. To distinguish E^+^H^+^ integrin molecules (Fig. 2k, bottom) from adjacent E^+^H^−^ and E^−^H^+^ integrins (Extended Data Fig. 6c), we compared the distributions of distances between KIM127 signals and mAb24 signals emanating from the same (filled gray curve) versus adjacent integrin molecules (open black curve) derived from simulations (Fig. 3a). The cross-over point between the two distributions was found to be at 8.2 nm. Thus, we considered all KIM127 signals within 8.2 nm of a mAb24 signal as evidence of a pair, i.e., an E^+^H^+^ integrin. We found about 4,600 E^+^H^+^ pairs on the microvillus tips within 50 nm from the coverslip. The E^+^H^+^ pairs accounted for 42% of all H^+^ and 46% of all E^+^ integrin molecules. This number was determined for each of 40 microvilli within 50 nm from the coverslip and showed 116±276 E^+^H^+^ integrin molecules per microvillus (Supplemental Table 2). According to our numerical simulations (made in Comsol)^31^ at a shear stress of 6 dyn/cm^2^, the drag force on a neutrophil with ~9.9 μm diameter (acquired from bright-field imaging of human neutrophils) is ~430 pN Therefore, at the time of neutrophil arrest, each microvillus provides an average of ~10.8 pN resisting force and each E^+^H^+^ integrin molecule supports a load of ~0.09 pN

**Fig. 3.**
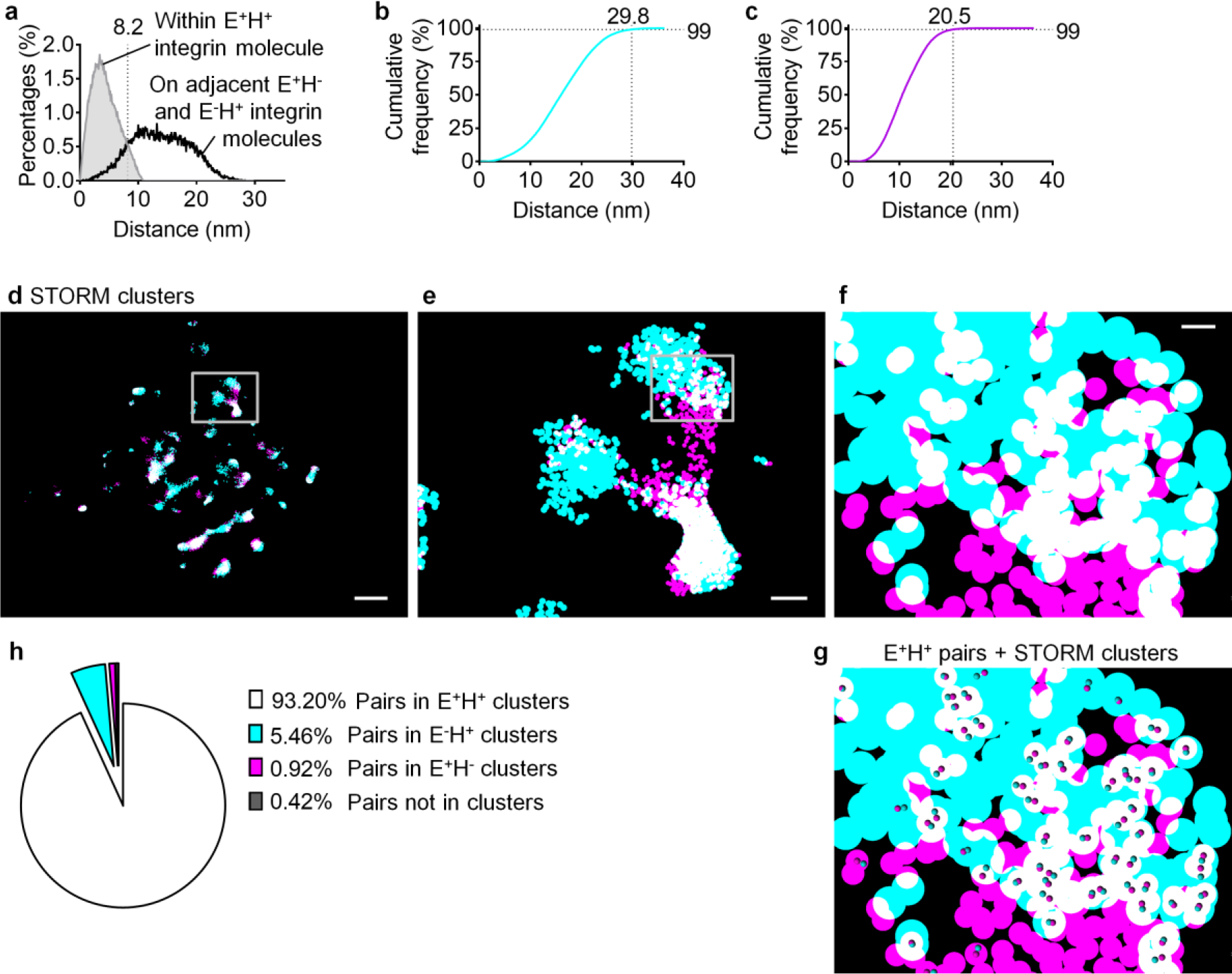
Molecular modeling defines the colocalization of E^+^ and H^+^ and clusters of E^+^H^−^, E^−^ H^+^ and E^+^H^+^ β_2_ integrins in the footprint of arrested neutrophils by STORM imaging. **a** To find E^+^H^+^ β_2_ integrin molecules with both mAb24 and KIM127 Fabs bound, distances between fluorophores on the same molecules were modeled (gray curves with filled area, 100,000 iterations), and compared with distances from adjacent E^−^H^+^ and E^+^H^−^ integrins (black curves with open area). The cutoff (crosses of the distributions) was 8.2 nm for pairs (KIM127 and mAb24 Fabs on same integrin molecule). **b**-**c** Clusters of mAb24 and KIM127 are defined based on distances of blinks obtained by random positioning of two adjacent integrins of the same type (64,000 iterations). Cumulative frequency of the randomly-simulated distances of mAb24 (**b**) or KIM127 (**c**) events. To cover 99% of clustered integrins, a cut-off of 29.8 or 20.5 nm for mAb24 and KIM127 was determined, respectively. **d**-**f** Binary cluster image of β_2_ integrin H^+^ (mAb24, cyan) and E^+^ (KIM127 Fab, magenta) on the footprint of a typical arrested human neutrophil. Overlap between magenta and cyan shown as white. Zoomed-in images in **e** (box in **d**) and **f** (box in **e**). **g** The E^+^H^+^ pairs (based on cutoff defined in **a**) were overlaid on the binary cluster image in **f**. Scale bars are 1μm for **d**, 200nm for **e**, and 30nm for **f**-**g**. **h** Frequency of the E^+^H^+^ pairs in each of the binarized cluster types. Data acquired from three neutrophil footprints.

Because integrins effectively operate as adhesion molecules localized in clusters^8,14^, we next determined the sizes and shapes of clusters. Existing clustering algorithms^32,33^ require arbitrary cutoffs in molecular density or distance. Since we know the shape of the projection of integrin molecules, we were able to construct cumulative histograms of the distances between KIM127 Fabs and mAb24 Fabs (Fig. 3b-c) without such arbitrary assumptions, just based on modeling integrins densely (“touching”) clustered in random orientations (Extended Data Fig. 6a-b). Using this approach, we found that 99% of mAb24 Fabs on randomly oriented, adjacent “touching” integrin molecules were within 29.8 nm of each other (Fig. 3b) and 99% of KIM127 Fabs on adjacent integrin molecules were within 20.5 nm of each other (Fig. 3c). Next, clusters of mAb24^+^ (H^+^) integrins were defined by merging circles with a radius of 29.8 nm around each mAb24 signal, and clusters of KIM127^+^ (E^+^) integrins were defined by merging circles with a radius of 20.5 nm around each KIM127 signal. This resulted in clearly defined clusters of various sizes and shapes for both E^+^ and H^+^ integrins (Fig. 3d-f, Extended Data Fig. 8a-b). The STORM clusters show a higher degree of ellipticity, elongated and even branched shapes, significantly different from the shapes of clusters visible in TIRF microscopy (Extended Data Fig. 8a-d). TIRF microscopy distorted the clusters to appear much more rounded (ellipticity closer to 1) because of insufficient resolution. Clusters of mAb24 were significantly larger than KIM127 clusters (p<0.0001, Extended Data Fig. 8e). STORM imaging identified significantly more (Extended Data Fig. 8h-i) but significantly smaller (Extended Data Fig. 8f-g, g-k) clusters than traditional TIRF imaging. In STORM imaging, most of the mAb24^+^ (~94%) and KIM127^+^ (~83%) events were clustered (Extended Data Fig. 8l).

We next colored H^+^ clusters cyan and E^+^ clusters magenta. Where H^+^ clusters overlapped with E^+^ clusters, we assigned white (Fig. 3d-f). The comparison of the colocalization area (overlap of H^+^ and E^+^ clusters) in STORM and TIRF shows that TIRF imaging overestimated the colocalization (Extended Data Fig. 8m). Since we knew the location of the E^+^H^+^ pairs, we next asked how the location of these pairs was related to the white clusters (Fig. 3g). About 93.2% of all E^+^H^+^ pairs were located within white clusters, ~5.5% within cyan, ~0.9% within magenta and ~0.4% were not located in clusters (isolated E^+^H^+^ integrins, Fig. 3h).

E^+^H^−^, E^−^H^+^ and E^+^H^+^ integrins might be oriented randomly within the clusters. To test this hypothesis, we modeled the distances between mAb24 Fabs and KIM127 Fabs in randomly distributed E^−^H^+^, E^+^H^−^, and E^+^H^+^ integrins (Fig. 4a-c). The modeled distribution for E^−^H^+^ integrins was compared to the measured distributions of distances in areas of the image where the resolution (σ) was better than 7.3 nm (Fig. 4d and data not shown). The measured distance distribution did not match the prediction from the random simulation. Thus, we conclude that E^+^H^−^ integrins are not randomly oriented on the microvilli of neutrophils. We next compared the modeled distance distribution to the distance distribution of E-H^+^ integrins in the entire footprint within 50 nm from the cover slip (Extended data figure 9) and found very similar results. Applying the same approach to E^+^H^−^ and E^+^H^+^ integrins showed that their distance distributions also deviated from random (Figure 4 e, f). For E^−^H^+^, E^+^H^−^ and E^+^H^+^ integrins, the measured distances were closer than those predicted by the random orientation model. In an attempt to find orientations consistent with the measured distance distributions, we considered “Face-to-Face” (integrins facing each other with the side where the Fab fragments are bound, Fig. 4j), “Back-to-Back” (integrins facing each other with the Fab fragments pointing away, data not shown) and “Parallel” orientations (integrins aligned with each other, Fig. 4k and l). The face-to-face orientation matched the experimental distance distribution of E^+^H^+^ (Fig. 4g), and parallel orientations matched the E^+^H^−^ (Fig. 4h) and E^−^H^+^ clusters (Fig. i).

That activated integrins are not oriented randomly in the clusters is an important new finding that likely impacts neutrophil arrest. Although our modeling excludes random orientations, we cannot be sure that the “Face-to-Face” and “Parallel” orientations are the true orientations. There is no unique solution, and other possible patterns might also fit the observed distance distributions. The “Face-to-Face” orientation of E^−^H^+^ integrins is supported by the known binding of E^−^H^+^ integrins to dimeric ICAM-1 and ICAM-3 in cis^8^ (Extended Data Fig. 10a). It is possible that E^−^H^+^ integrins are forced to face each other, on average, because many of them may be bound to the same ICAM dimer (Extended Data Fig. 10b). Kindlin-3 is critical for integrin activation^34–36^ and relevant to integrin clustering^14^. A recent study showed that kindlin forms dimers^37^, which could also participate in setting up the “Face-to-Face” pattern of E^−^H^+^ integrins. In conclusion, we show the first map of activated integrins during cell arrest. This work defines the number of activated integrins needed to arrest a rolling neutrophil and refines their positions, cluster size and shape. In addition, our data reveal an unexpected non-random patterning of activated β_2_ integrins.

**Fig. 4.**
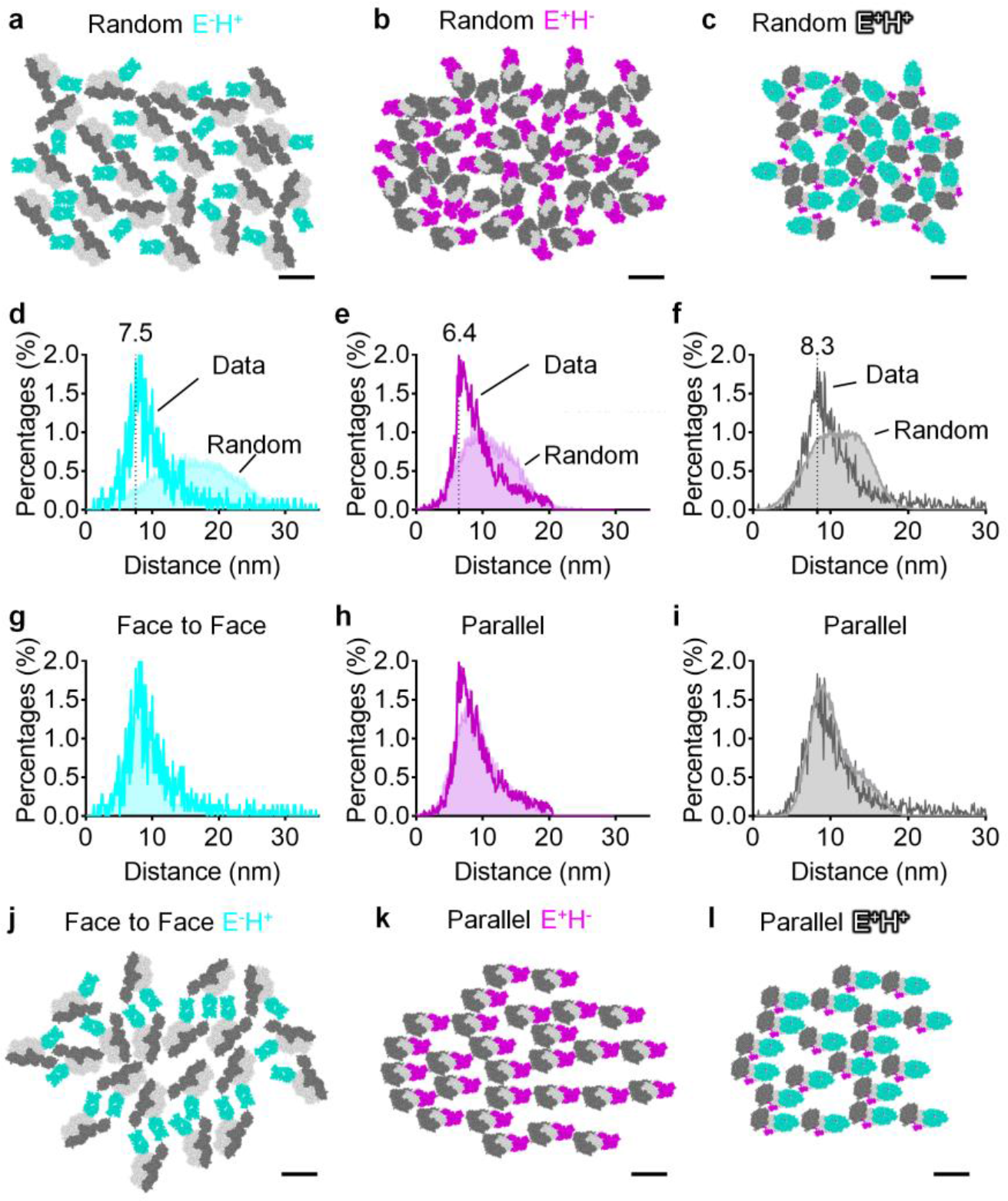
Molecular pattern of β_2_ integrin activation. **a**-**c** Typical space-filling top views of E^−^H^+^ (**a**), E^+^H^−^ (**b**), and E^+^H^+^ (**c**) β_2_ integrins with random directions and orientations. **d**-**i** Distance distributions of E^−^H^+^ (σ≤7.3nm, **d**, **g**), E^+^H^−^ (**e**, **h**) and E^+^H^+^ (**f**, **i**) integrins in STORM images within 50nm to the coverslip (dark cyan, magenta, or grey curves with open area, respectively) compared with distance distributions from random simulations (light cyan, magenta, or grey curves with filled area, respectively). The random simulations in **d**-**f** contain 64 conditions (8 directions × 8 orientations, 64,000 iterations each). The random simulations in **g** contain sixteen‘Face-to-Face’ conditions. The random simulations in **h** and **i** contain eight ‘Parallel’ conditions each. **j**-**1** Examples showing space-filling E^−^H^+^ (**j**) integrins ‘Face-to-Face’, E^+^H^−^ (**k**) and E^+^H^+^ (**1**) integrins ‘Parallel’, which best fit the observed distance distribution. Scale bars are all 10nm.

## Methods

**Reagents.** Recombinant human P-selectin-Fc, ICAM-1-Fc, and IL-8 were purchased from R&D Systems. Casein blocking buffer, fluorescein (FITC), and CellTracker Orange CMRA were purchased from Thermo Fisher Scientific. The Fab fragments of the conformation-specific antibody, mAb24, to human β_2_-I-like-domain, which reports the headpiece-opening ^12,13^, were contract-manufactured by Biolegend. The KIM127 mAb to human β_2_-IEGF-domain, which reports the ectodomain extension^10,11^, was purified at the Lymphocyte Culture Center at the University of Virginia from hybridoma supernatant (ATCC). The Fab fragments of KIM127 were produced by the Pierce™ Fab Preparation Kit from Thermo Fisher Scientific. The Fabs of mAb24 and KIM127 were directly labeled by DL550 or DL650 using DyLight antibody labeling kits from Thermo Fisher Scientific. Alexa Fluor 488 (AF488) conjugated mouse anti-human CD16 monoclonal antibody was purchased from Biolegend. Quantum Simply Cellular anti-Mouse (calibration beads) was purchased from Bangs Laboratories. Polymorphprep was purchased from Accurate Chemical. Roswell Park Memorial Institute (RPMI) medium 1640 without phenol red and phosphate-buffered saline (PBS) without Ca^2+^ and Mg^2+^ were purchased from Gibco. Human Serum Albumin (HSA) was purchased from Gemini Bio Products. Paraformaldehyde (PFA) was purchased from Thermo Fisher Scientific. The STORM buffer contains 50 mM Tris pH 8.0, 10 mM NaCl, 10 % Glucose, 0.1 M Mercaptoethanolamine (Cysteamine Sigma-Aldrich), 56 U/ml Glucose Oxidase (from Aspergillus niger, Sigma-Aldrich), and 340 U/ml Catalase (from bovine liver, Sigma-Aldrich).

**Neutrophil isolation.** Heparinized whole blood was obtained from healthy human donors after informed consent, as approved by the Institutional Review Board of the La Jolla Institute of Allergy & Immunology in accordance with the Declaration of Helsinki. Neutrophils were isolated by using Polymorphprep (a mixture of sodium metrizoate and Dextran 500) density gradient (Accurate Chemical)^8^. Briefly, human blood was applied onto Polymorphprep, centrifuged at 500 g for 35 min at 20-25°C, resulting in neutrophils concentrated in a layer between peripheral blood mononuclear cells and erythrocytes. After washing with PBS without Ca^2+^ and Mg^2+^ twice, the neutrophils (>95% purity by flow cytometry, no visible activation by microscopy) were re-suspended in RPMI-1640 without phenol red plus 2% HSA and were used within four hours. Neutrophils were incubated with FcR blocking reagents for ten minutes at room temperature (RT) prior to all the experiments.

**Microfluidic device.** The assembly of the microfluidic devices used in this study and the coating of coverslips with recombinant human P-selectin-Fc and ICAM-1-Fc has been described previously^8,18,21,38^. Briefly, cleaned coverslips were coated with P-selectin-Fc (2 μg ml^−1^) and ICAM-1-Fc (10 μg ml^−1^) for 2 hours and then blocked for 1 hour with casein (1%) at RT. After coating, coverslips were sealed to polydimethylsiloxane (PDMS) chips by magnetic clamps to create flow chamber channels ~29 μm high and ~300 μm across. By modulating the pressure between the inlet well and the outlet reservoir, 6 dyn cm^−2^ wall shear stress was applied in all experiments.

**Microfluidic perfusion assay.** To fix the neutrophils at the time of arrest, the time of arrest upon IL-8 and the time of fixation upon PFA were assessed (Extended Data Fig. 1). Isolated human primary neutrophils (5×10^6^ cells ml^−1^) were labeled with 10 μM CellTracker Orange CMRA at RT for 10 minutes and perfused in the microfluidic device over a substrate of recombinant human P-selectin-Fc and recombinant human ICAM-1-Fc at a shear stress of 6 dyn cm^−2^. IL-8 (10 ng ml^−1^) or PFA (8%) was mixed with FITC (1μM) and perfused through the microfluidic flow chamber after cell rolling on the substrate. Dual-color epifluorescence imaging (CMRA and FITC) was recorded by an IX71 inverted research microscope (Olympus America) with a 40× air objective to monitor the arrival of IL-8 or PFA by FITC and the stop (arresting by IL-8 or fixing by PFA) of neutrophils. The background mean fluorescence intensity (MFI) of FITC and the cell velocity was quantified by using FIJI-ImageJ2^39^. Cell arrest was defined as the time when the velocity dropped below 0.1 μm s^−1^. IL-8 was added 12 seconds before PFA to achieve fixation at the moment of arrest (Extended Data Fig. 1).

To prepare for STORM microscopy, isolated human primary neutrophils (5×10^6^ cells ml^−1^) were incubated with fluorochrome-conjugated reporting Fabs (mAb24-DL550 and KIM127-DL650, 5 μg ml^−1^ each) for 3 minutes at RT and immediately perfused through the microfluidic device over a substrate of recombinant human P-selectin-Fc and recombinant human ICAM-1-Fc at a wall shear stress of 6 dyn cm^−2^ without separation of the unbound Fabs. After cells began rolling on the substrate, 10 ng ml^−1^ IL-8 was perfused for twelve seconds, then 8% PFA was perfused for five minutes to fix the cells at the time of arrest. After washing with the medium (RPMI-1640 without phenol red + 2% HSA) for five minutes, Fabs (mAb24-DL550 and KIM127-DL650, 5 μg ml^−1^ each) were perfused again to saturate the staining of active epitopes. Coverslips were disassembled from the microfluidic device and mounted with the STORM buffer on a glass slide prior to imaging.

**STORM setup and imaging.** Images were captured using 100× 1.49 NA Apo TIRF objective with TIRF (80° incident angle) illumination on a Nikon Ti super-resolution microscope. Images were collected on an ANDOR IXON3 Ultra DU897 EMCCD camera using the multicolor sequential mode setting in the NIS-Elements AR software (Nikon Instruments Inc., NY). Power on the 488, 561, and 647-nm lasers was adjusted to 50% to enable collection of between 100 and 300 blinks per 256 × 256 pixel camera frame in the center of the field at appropriate threshold settings for each channel. The collection was set to 20,000 frames, yielding 1-2 million molecules.

**Molecular Modeling.** CCP4MG (http://www.ccp4.ac.uk/MG) is used in structural modeling and generating figures^40^. We used the published structure of E^−^H^+^ β_2_ integrin extracellular domains (PDB: 4NEH)^27^ and modeled a Fab (PDB: 3HI6) on the binding site of mAb24^13^. For the structure of E^+^H^−^ β_2_ integrin, published E^−^H^−^ β_2_ integrin extracellular domains (PDB: 3K6S)^26^ were adjusted to model an extended headpiece. Modeled structures were matched to EM images of E^+^ β_2_ integrin^24^. A Fab was modeled to the binding site of KIM127^11^. For the structure of E^+^H^+^ β_2_ integrin, the hybrid domain swing out of E^−^H^+^ β_2_ integrin (PDB: 4NEH) was superposed with the open headpiece of integrin α_IIB_β_3_ (PDB: 3FCU)^28^. To match EM images of E^+^H^+^ β_2_ integrin^25^, the headpiece was unbent, the membrane-proximal ends of integrin α and β chains were modeled close to each other, and the membrane-distal ends of integrin α and β tailpieces were modeled linked to the membrane-proximal ends of integrin α and β headpieces. Two Fabs were modeled to the binding sites of mAb24^13^ and KIM127^11^. Two-dimensional STORM data were acquired and the distances of bound Fabs were estimated in the top view projection.

**Image processing – generation of STORM pointillisms.** The super-resolution localizations were reconstructed with the Nikon STORM software^41^. Positions of individual blinks have been localized with high accuracy by switching them on and off sequentially using the 488, 561, and 647-nm lasers at appropriate power settings. Fitting the histograms of beads with Gaussians gave standard deviations of 17.25 nm and 15.88 nm for the DL550 channel in the x and y directions, respectively. Analogous values for the DL650 channel were 15.68 nm and 14.97 nm (Extended Data Fig. 2a). The positions determined from multiple switching cycles can show a substantial drift over the duration of the acquisition. This error is considerably reduced by correcting for sample drift over the course of the experiment by an auto-correlation method reconstructed from 200-1000 frames (implemented in the Nikon software)^41^. The number of frames used in a set is based on the number of blinks identified, with a default setting of 10,000 blinks. Displacement is corrected by translation and rotation in the X and Y directions for 2D STORM. Axial drift over the course of the acquisition is minimized by engaging the Nikon Perfect Focus System. Calibration of chromatic shift (warp correction) was carried out using a multicolored 100nm TetraSpeck beads using at least 100 beads per field. Calibration for warp correction for 2D STORM was executed using the 2D warp calibration feature of the Nikon STORM software. Briefly, a total of 201 images were collected for each channel (488, 561 and 647nm) without the cylindrical lens in place. Frames 1-20 and frames 182-201 are collected at the focal position. Frames 21-181 are collected across a range of 1.6 microns in 10 nm steps in the Z (covering 800 nm above and 800 nm below the focal plane). The calibration files generated from this software feature were applied during analysis for the correction of the STORM images. Blinking events were followed for successive frames to filter out blinks generating traces longer than five frames during analysis. Furthermore, only molecules with a point spread function (PSF) of 200-400nm (based on a 100× 1.49 objective) and a photon count above 100 (based on camera noise of the ANDOR EMCCD) were retained. The data was further filtered based on empirical observation of photon count signals (peak height when converted to an intensity value) found in cells vs background staining on the glass slide surface (generally values above 300-700 intensity units above camera noise). The precision of the localization during a switching cycle is calculated from these parameters and from photon counts using molecules that are ultimately well separated in the sample itself^15,41,42^. After getting the datasheet of the blink events, Imaris was used to generate the pointillism map (Fig. 1i-k).

**Image processing – filtering the blinks within the reach of the substrate.** Neutrophils have microvilli that are up to ~200 nm high^20^ and the sum of the length of an extended β_2_ integrin and its ligand ICAM-1-Fc is ~50 nm^8,17^. Only integrins on the “tops” of microvilli can reach (and bind) ICAM-1, and other integrins on the “sides” of the microvilli or in the “valleys” between the microvilli are too far away from the adhesive substrate to contribute to binding^8^, even though such integrin molecules may be in the fully activated E^+^H^+^ conformation that could bind ligand in trans if they could reach it (Extended Data Fig. 4a). Only β_2_ integrins that could physically bind ICAM-1 in trans can contribute to arrest. Since the intensity of the TIRF signal emanating from the homogeneous membrane label (CD16, a Glycosylphosphatidylinositol (GPI)-anchored protein) mono-exponentially decays with the distance from the substrate^8,18,21,22^, we used CD16-AF488 intensity to construct a three-dimensional (3D) “hills and valleys” plot. We filtered for mAb24 and KIM127 Fab signals that were in areas in which the CF16 signal was within 50 nm of the coverslip (Fig. 2a, Extended Data Fig. 4, Matlab codes available at Github). By using such a narrow slice, we select only the “tips” of microvilli which form planes nearly parallel to the coverslips (Extended Data Fig. 4). Integrin molecules anchored in the membrane plane are approximately normal to the coverslip, which guided our top-view molecular modeling (see Supplementary Movies 1-3). Imaris was used to generate the pointillism maps after the proximity filtering (Fig. 2c, d, n, o).

**Image processing – eliminating multiple blinks from each Fab.** Because multiple blinks of the same fluorochrome may be recorded during STORM imaging, and one Fab may be labeled with several fluorochromes, it is necessary to merge these blinks to one Fab event for further analysis. We used molecular modeling to find which blinks likely represent the same Fab molecule. First, we modeled the structure of various β_2_ integrins with Fab molecules attached as described above.

To find the cut-off for merging blinks, fluorochromes were placed randomly in each Fab, and the projection of their distance in the plane of the coverslip was recorded. 100,000 simulations generated the distribution of the distance of the fluorochromes residing in the same Fab (Fig. 2i, m, the curve with dark cyan and magenta line and open area).

This was repeated for blinks emanating from two fluorochromes located in Fabs bound to two different but adjacent integrin molecules. In each simulation, integrins are closely packed (allowed to touch), but randomly oriented. We tested eight positions of the second integrin molecule relative to the first integrin molecule (top, bottom, left, right, top-left, top-right, bottom-left, bottom-right) and eight orientations (one of the integrins was rotated by 0°, 45°, 90°, 135°, 180°, 225°, 270°, 315° about the z axis), totaling 64 conditions for the relative position of the two integrins. One thousand random simulations were done for each of these conditions. These 64,000 simulations yielded the distribution of the distance of blinks emanating from fluorochromes in two Fabs bound to adjacent integrins (Fig. 2i, m, the curve with light cyan and magenta line and closed area). The cross of the two distributions was used as the cut-off for merging. 99.1% or 96.8% of blinks truly emanating from the same mAb24 or KIM127 Fab, respectively, and 7.8% or 11.5% of blinks emanating from mAb24 or KIM127 Fabs, respectively, bound to adjacent integrin molecules were correctly assigned. ThunderSTORM^43^ was used to eliminate the multiple blinks in the same Fab within the calculated distance threshold. Imaris was used to generate the pointillism map after eliminating multiple blinks (Fig. 2p).

**Image processing – colocalization.** To obtain the molecular cut-off for colocalized pairs, i.e., the E^+^H^+^ integrin molecules with both mAb24 and KIM127 Fabs bound, two new random simulations were performed. If a KIM127 and a mAb24 Fab are bound to the same E^+^H^+^ integrin, the blinks emanating from their fluorochromes were assigned the range of possible positions in the projection of the coverslip plane. In each simulation, a fluorochrome is randomly placed in the KIM127 Fab and another in the mAb24 Fab, and the distances are calculated. 100,0 such simulations yielded the distribution of the distances of blinks emanating from fluorochromes in KIM127 and mAb24 Fabs on the same E^+^H^+^ integrin (Fig. 3a, the curve with gray line and closed area). For KIM127 and mAb24 Fabs are bound to adjacent E^+^H^−^ and E^−^H^+^ integrin molecules, a second distance distribution is generated. Similar to the approach used above, the integrin molecules (E^+^H^−^ and E^−^H^+^) are closely packed and randomly positioned and oriented, totaling 64 conditions for the relative position of the two integrins. 64,000 random simulations (1,000 for each position and orientation) yield the distribution of the distances of blinks emanating from KIM127 and mAb24 Fabs bound to adjacent E^+^H^−^ and E^−^H^+^ integrins (Fig. 3a, the curve with black line and open area). The cross of the two distributions was used as the cut-off for merging. 93.8% of KIM127-mAb24 pairs truly emanating from the same integrin and 14.4% of KIM127-mAb24 pairs from Fabs bound to adjacent integrin molecules were correctly assigned. KIM127-mAb24 pairs were identified with a custom Matlab algorithm (available at Github). Briefly, KIM127 and mAb24 mutual nearest neighbor pairs (where a KIM127^+^ molecule and a mAb24^+^ integrin molecule identify each other as their nearest neighbors of the other species) were found using“KdTreeSearcher” in Matlab. These pairs were filtered from the list of H^+^ and E^+^ positions, and then the nearest neighbor search was repeated until no new pairs were identified. Only those pairs with a separation distance less than the calculated threshold (8.2 nm) were kept for the final calculation. Imaris was used to generate the image of colocalization pairs (Fig. 2j).

**Image processing – clustering.** To define clusters, we constructed circles around the most likely center of each KIM127 and each mAb24 signal with radii (20.5 and 29.8 nm, respectively) that enclosed 99% of all distances between adjacent integrins from the random simulations (Fig. 3b, c). The integrins in circles touching or overlapping a circle of its same kind (KIM127^+^ or mAb24^+^, respectively) were considered to be in clusters. Thus, a cluster could be as small as two and as large as thousands of integrin molecules. These STORM-based binary clusters (Fig. 3d-f) were generated by masking from the pointillism map (with the diameters of the above clustering cut-off) after eliminating multiple blinks (with the diameter as the cluster cut-off) in Imaris. Sizes, ellipticities, number of clusters and cluster areas (Extended Data Fig. 8) were obtained in Imaris.

**Statistics.** Statistical analysis was performed with Prism 6 (GraphPad). Data are presented as violin plots (Extended Data Fig. 8c, d), bean plots (Extended Data Fig. 8f, g)^44^, mean ± SD (Extended Data Fig. 1a-b, Fig. 8h-m), distributions (Fig. 2b, e-f, l-m, 3 a, 4d-I, Extended Data Fig. 6, Extended Data Fig. 2a, Extended Data Fig. 8e) and cumulative distributions (Fig. 3b-c), pie chart (Fig. 3h). The means for the data sets were compared using paired student t-tests with equal variances (Extended Data Fig. 8h-m) or Mann-Whitney test (Extended Data Fig. 8c-g). P values less than 0.05 were considered significant. Post hoc analysis (SPSS 22.0 software, IBM) was performed to ensure that the sample size we chose had adequate power.

## Data availability

The data that support the findings of this study are available from the corresponding author upon request. The raw data sets are large (gigabytes each) and require special data transfer arrangements.

## Acknowledgements

This research was supported by funding from the National Institutes of Health, USA (NIH, HL078784), WSA postdoctoral fellowship from the American Heart Association, USA (AHA, 16POST31160014), and Career Development Award from the AHA (18CDA34110426). We acknowledge Dr. Sara McArdle from the Microscopy Core Facility, La Jolla Institute for Allergy and Immunology for coding the custom scripts on MatLab used in this study. We acknowledge Dr. Yunmin Jung from the Division of Inflammation Biology, La Jolla Institute for Allergy and Immunology for assessing the resolution of the STORM microscope. We acknowledge both Dr. Sara McArdle and Dr. Zbigniew Mikulski from the Microscopy Core Facility, La Jolla Institute for Allergy and Immunology for providing help of imaging processing. We acknowledge Dr. Kersi Pestonjamasp from the Moores Cancer Center, University of California San Diego for the maintaining and calibration of the STORM microscope, and helping us with the data acquisition and analysis.

## Author Contributions

Experiments were designed by Z.F. and K.L. Most experiments were performed by Z.F. and W.B.K. Structural modeling is performed by Z.F. and assessed by D.M.Z and M.A.A. Image processing was performed by Z.F. and W.B.K. Data analysis was performed by Z.F. The microfluidic device was designed by E.G. and A.G. The simulation of neutrophil load bearing was performed by A.G.The manuscript was written by K.L., Z.F. and W.B.K. The project was supervised by K.L. All authors discussed the results and commented on the manuscript.

## Author Information

The authors declare no competing financial interest. Correspondence and requests for materials should be addressed to.

## References

1 Arfors, K.-E. et al. A monoclonal antibody to the membrane glycoprotein complex CD18 inhibits polymorphonuclear leukocyte accumulation and plasma leakage in vivo. Blood 69, 338–340 (1987).

2 von Andrian, U.H. et al. Two step model of leukocyte-endothelial cell interaction in inflammation: Distinct roles for LECAM-1 and the leukocyte b_2_ integrins in vivo. Proc Natl Acad Sci USA 88, 7538–7542 (1991).

3 Lawrence, M.B. & Springer, T.A. Leukocytes roll on a selectin at physiologic flow rates: Distinction from and prerequisite for adhesion through integrins. Cell 65, 859–873 (1991).

4 Xiong, J.P. et al. Crystal structure of the extracellular segment of integrin alpha Vbeta3. Science 294, 339–345 (2001).

5 Shimaoka, M. et al. Stabilizing the integrin alpha M inserted domain in alternative conformations with a range of engineered disulfide bonds. Proc. Natl. Acad. Sci. U. S. A 99, 16737–16741 (2002).

6 Shimaoka, M., Takagi, J. & Springer, T.A. Conformational regulation of integrin structure and function. Annual Review of Biophysics & Biomolecular Structure 31, 485–516 (2002).

7 Takagi, J., Petre, B., Walz, T. & Springer, T. Global conformational rearrangements in integrin extracellular domains in outside-in and inside-out signaling. Cell 110, 599–611 (2002).

8 Fan, Z. et al. Neutrophil recruitment limited by high-affinity bent beta2 integrin binding ligand in cis. Nat Commun 7, 12658 (2016).

9 Sundd, P. et al. Dynamic footprinting reveals mechanisms of leukocyte rolling. Nature Methods 7, 821–824 (2010).

10 Robinson, M.K. et al. Antibody against the Leu-CAM beta-chain (CD18) promotes both LFA-1- and CR3-dependent adhesion events. J Immunol 148, 1080–1085 (1992).

11 Lu, C., Ferzly, M., Takagi, J. & Springer, T.A. Epitope Mapping of Antibodies to the C-Terminal Region of the Integrin beta2 Subunit Reveals Regions that Become Exposed Upon Receptor Activation. The Journal of Immunology 166, 5629–5637 (2001).

12 Dransfield, I. & Hogg, N. Regulated expression of Mg2+ binding epitope on leukocyte integrin alpha subunits. EMBO J 8, 3759–3765 (1989).

13 Yang, W., Shimaoka, M., Chen, J. & Springer, T.A. Activation of integrin beta-subunit I-like domains by one-turn C-terminal alpha-helix deletions. Proc Natl Acad Sci U S A 101, 2333–2338 (2004).

14 Ye, F. et al. The mechanism of kindlin-mediated activation of integrin alphaIIbbeta3. Curr Biol 23, 2288–2295 (2013).

15 Rust, M.J., Bates, M. & Zhuang, X. Sub-diffraction-limit imaging by stochastic optical reconstruction microscopy (STORM). Nat Methods 3, 793–795 (2006).

16 Betzig, E. et al. Imaging intracellular fluorescent proteins at nanometer resolution. Science 313, 1642–1645 (2006).

17 Moore, T.I., Aaron, J., Chew, T.L. & Springer, T.A. Measuring Integrin Conformational Change on the Cell Surface with Super-Resolution Microscopy. Cell Rep 22, 1903–1912 (2018).

18 Sundd, P. et al. Quantitative dynamic footprinting microscopy reveals mechanisms of neutrophil rolling. Nat Methods 7, 821–824 (2010).

19 Bon, P. et al. Three-dimensional nanometre localization of nanoparticles to enhance super-resolution microscopy. Nat Commun 6, 7764 (2015).

20 Bruehl, R.E., Springer, T.A. & Bainton, D.F. Quantitation of L-selectin distribution on human leukocyte microvilli by immunogold labeling and electron microscopy. J Histochem Cytochem 44, 835–844 (1996).

21 Sundd, P. et al. ’slings’ enable neutrophil rolling at high shear. Nature 488, 399–403 (2012).

22 Jung, Y. et al. Three-dimensional localization of T-cell receptors in relation to microvilli using a combination of superresolution microscopies. Proc Natl Acad Sci U S A 113, E5916–E5924 (2016).

23 Zhang, H. et al. Structural basis of activation-dependent binding of ligand-mimetic antibody AL-57 to integrin LFA-1. Proc Natl Acad Sci U S A 106, 18345–18350 (2009).

24 Chen, X. et al. Requirement of open headpiece conformation for activation of leukocyte integrin alphaXbeta2. Proc Natl Acad Sci U S A 107, 14727–14732 (2010).

25 Chen, X., Yu, Y., Mi, L.Z., Walz, T. & Springer, T.A. Molecular basis for complement recognition by integrin alphaXbeta2. Proc Natl Acad Sci U S A 109, 4586–4591 (2012).

26 Xie, C. et al. Structure of an integrin with an alphaI domain, complement receptor type 4 EMBO J29, 666–679 (2010).

27 Sen, M., Yuki, K. & Springer, T.A. An internal ligand-bound, metastable state of a leukocyte integrin, alphaXbeta2. J Cell Biol 203, 629–642 (2013).

28 Zhu, J. et al. Structure of a complete integrin ectodomain in a physiologic resting state and activation and deactivation by applied forces. Mol Cell32, 849–861 (2008).

29 Sengupta, P., Jovanovic-Talisman, T. & Lippincott-Schwartz, J. Quantifying spatial organization in point-localization superresolution images using pair correlation analysis. Nat Protoc 8, 345–354 (2013).

30 Alon, R. & Feigelson, S.W. Chemokine-triggered leukocyte arrest: force-regulated bi-directional integrin activation in quantal adhesive contacts. Curr Opin Cell Biol 24, 670–676 (2012).

31 Marki, A., Gutierrez, E., Mikulski, Z., Groisman, A. & Ley, K. Microfluidics-based side view flow chamber reveals tether-to-sling transition in rolling neutrophils. Sci Rep 6, 28870 (2016).

32 Levet, F. et al. SR-Tesseler: a method to segment and quantify localization-based super-resolution microscopy data. Nat Methods 12, 1065–1071 (2015).

33 Andronov, L., Orlov, I., Lutz, Y., Vonesch, J.L. & Klaholz, B.P. ClusterViSu, a method for clustering of protein complexes by Voronoi tessellation in super-resolution microscopy. Sci Rep 6, 24084 (2016).

34 Moser, M., Nieswandt, B., Ussar, S., Pozgajova, M. & Fassler, R. Kindlin-3 is essential for integrin activation and platelet aggregation. Nat Med 14, 325–330 (2008).

35 Moser, M. et al. Kindlin-3 is required for β_2_ integrin-mediated leukocyte adhesion to endothelial cells. Nat Med 15, 300–305 (2009).

36 Lefort, C.T. et al. Distinct roles for talin-1 and kindlin-3 in LFA-1 extension and affinity regulation. Blood 119, 4275–4282 (2012).

37 Li, H. et al. Structural basis of kindlin-mediated integrin recognition and activation. Proc Natl Acad Sci U S A 114, 9349–9354 (2017).

38 Sundd, P. et al. Live cell imaging of paxillin in rolling neutrophils by dual-color quantitative dynamic footprinting. Microcirculation 18, 361–372 (2011).

39 Rueden, C.T. et al. ImageJ2: ImageJ for the next generation of scientific image data. BMC Bioinformatics 18, 529 (2017).

40 McNicholas, S., Potterton, E., Wilson, K.S. & Noble, M.E. Presenting your structures: the CCP4mg molecular-graphics software. Acta Crystallogr D Biol Crystallogr 67, 386–394 (2011).

41 Huang, B., Wang, W., Bates, M. & Zhuang, X. Three-dimensional super-resolution imaging by stochastic optical reconstruction microscopy. Science 319, 810–813 (2008).

42 Thompson, R.E., Larson, D.R. & Webb, W.W. Precise nanometer localization analysis for individual fluorescent probes. Biophys J82, 2775–2783 (2002).

43 Ovesny, M., Krizek, P., Borkovec, J., Svindrych, Z. & Hagen, G.M. ThunderSTORM: a comprehensive ImageJ plug-in for PALM and STORM data analysis and super-resolution imaging. Bioinformatics 30, 2389–2390 (2014).

44 Kampstra, P. Beanplot: A Boxplot Alternative for Visual Comparison of Distributions. 2008 28, 9 (2008).

